# Natural genetic variation screen in *Drosophila* identifies Wnt signaling, mitochondrial metabolism, and redox homeostasis genes as modifiers of apoptosis

**DOI:** 10.1101/670075

**Authors:** Rebecca A.S. Palu, Elaine Ong, Kaitlyn Stevens, Shani Chung, Katie G. Owings, Alan G. Goodman, Clement Y. Chow

## Abstract

Apoptosis is the primary cause of degeneration in a number of neuronal, muscular, and metabolic disorders. These diseases are subject to a great deal of phenotypic heterogeneity in patient populations, primarily due to differences in genetic variation between individuals. This creates a barrier to effective diagnosis and treatment. Understanding how genetic variation influences apoptosis could lead to the development of new therapeutics and better personalized treatment approaches. In this study, we examine the impact of the natural genetic variation in the *Drosophila* Genetic Reference Panel (DGRP) on two models of apoptosis-induced retinal degeneration: overexpression of *p53* or *reaper* (*rpr*). We identify a number of known apoptotic, neural, and developmental genes as candidate modifiers of degeneration. We also use Gene Set Enrichment Analysis (GSEA) to identify pathways that harbor genetic variation that impact these apoptosis models, including Wnt signaling, mitochondrial metabolism, and redox homeostasis. Finally, we demonstrate that many of these candidates have a functional effect on apoptosis and degeneration. These studies provide a number of avenues for modifying genes and pathways of apoptosis-related disease.

## INTRODUCTION

Phenotypic heterogeneity is the driving force behind the Precision Medicine Initiative (Scriver and Waters 1999; Nadeau 2001; Queitsch *et al.* 2012; Gallati 2014). Patients suffering from the same genetic disorders can carry identical causal mutations but often display wildly variable phenotypes and symptom severity. A large part of this variation is due to inter-individual differences in genetic background, including silent cryptic genetic variation that is revealed upon disease or stress (Queitsch *et al.* 2012; Chow 2016). Understanding the role of this variation and the genes or pathways which modify disease will lead to improved personalized therapeutic predictions, strategies, and diagnostics.

One process implicated in many genetic disorders is programmed cell death or apoptosis (Elmore 2007; Sano and Reed 2013; Kurtishi *et al.* 2018). During normal development and tissue turnover, cells can receive both internal and external signals that trigger a programmed response which eventually results in the death of the cell (Elmore 2007). Because cell death is essential to cellular, tissue, and organismal homeostasis, disruption of apoptosis pathways can be catastrophic. Inhibition of apoptosis is an important step in transformation and cancer, while excess apoptosis, often activated by chronic cellular stress, is a primary cause of degeneration in different neuronal, retinal, muscular-skeletal, and metabolic diseases (Mattson 2000; Elmore 2007; Ouyang *et al.* 2012). As a result, an important area of therapeutic development is focused on targeting apoptosis without disrupting normal tissue homeostasis (Elmore 2007). Our previous work demonstrated that variation in apoptotic genes is associated with phenotypic variation in a model of retinal degeneration, suggesting that modifiers of apoptosis could serve as drug targets in degenerative diseases (Chow *et al.* 2016).

Model organism tools, such as the *Drosophila* Genetic Reference Panel (DGRP), enable the study of the impact of natural genetic variation on diseases and related pathways. The DGRP is a collection of ~200 isogenic strains derived from a wild population, such that each strain represents one wild-derived genome (Mackay *et al.* 2012). The variation in the DGRP is well tolerated under healthy, non-disease conditions and allows for the identification of genetic polymorphisms that are associated with phenotypic variation in models of human disease (Chow and Reiter 2017). Importantly, the availability of full-genome sequence for these strains allows for genome-wide association analyses that link quantitative phenotypes with genetic variation and modifier genes.

In this study, we report the results of natural variation screens of *reaper*-(*rpr*) and *p53*-induced apoptosis (Figure 1). Overexpression of either of these genes leads to massive apoptotic activation (Hay *et al.* 1995; Jin *et al.* 2000). While there is a great deal of overlap between these pathways, they can each activate apoptosis independently. p53 is stabilized in response to DNA damage and initiates apoptosis by transcriptionally activating the inhibitor of apoptosis (IAP) inhibitors *rpr*, *grim*, and *hid* (Mollereau and Ma 2014) (Figure 1). It also induces the expression of the *Drosophila* TNF *eiger*, which subsequently increases apoptosis by activating JNK signaling and stabilizing the IAP inhibitor Hid (Shklover *et al.* 2015). *rpr* is activated transcriptionally by either p53 or the JNK signaling cascade, which is induced downstream of oxidative, ER, and other cellular stresses (Kanda and Miura 2004; Shlevkov and Morata 2012) (Figure 1). We designed this study to identify genetic modifiers of general apoptosis as well as modifiers that are specific to stress-induced, p53-independent pathways.

**Figure 1.**
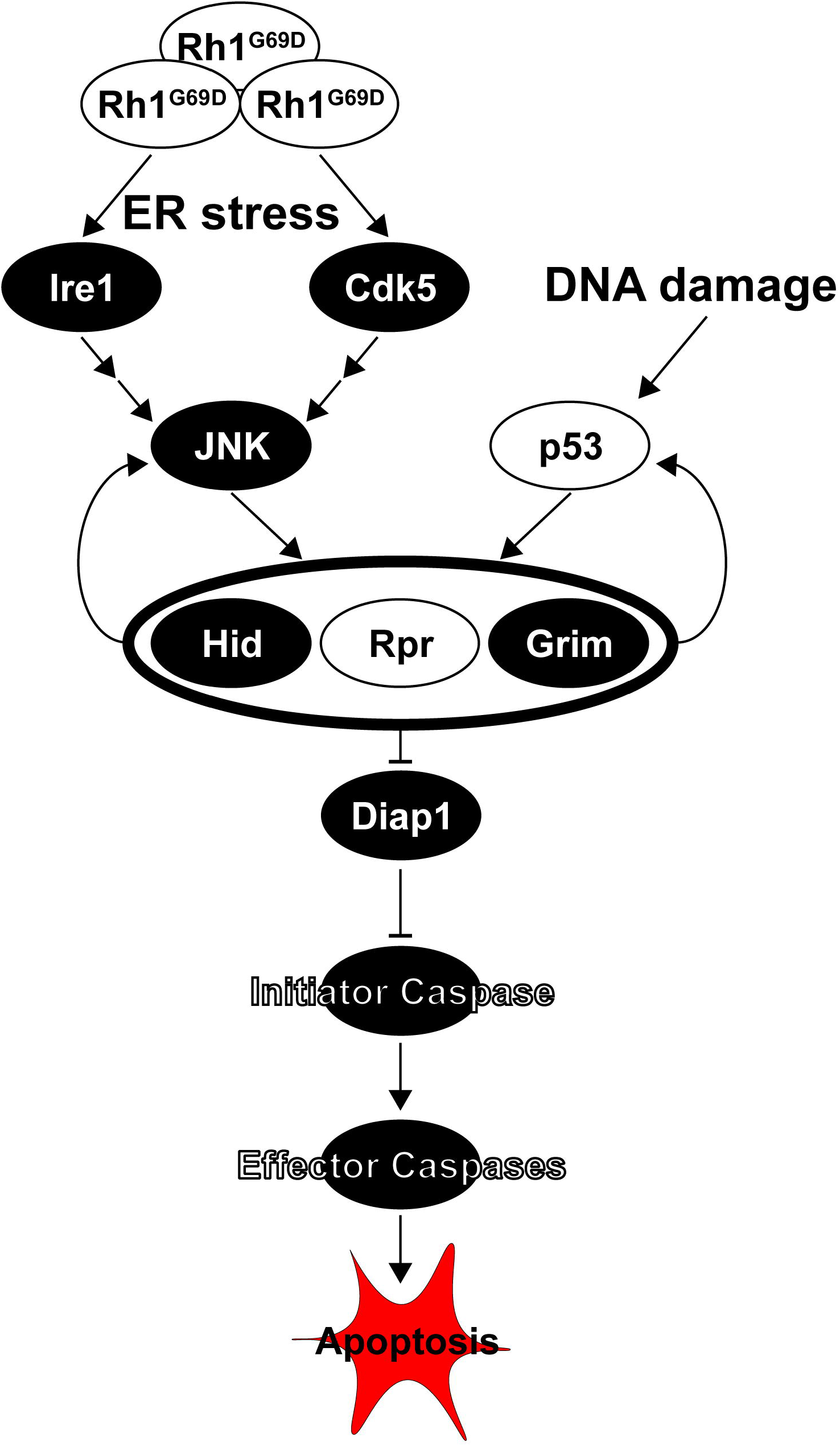
Activation of apoptosis through *p53* and *rpr*-associated pathways. Apoptosis is primarily initiated through either p53 or Jun-induced (JNK) transcriptional activation of the Inhibitor of Apoptosis (IAP, in *Drosophila* Diap) inhibitors *hid*, *rpr* and *grim*. While p53 is primarily activated by DNA damage and disruption of the cell cycle, JNK signaling is activated downstream of cellular stress, such as endoplasmic reticulum (ER) stress, through Ire1 and Cdk5. ER stress occurs when misfolding proteins, like the rhodopsin mutant *Rh1*^*G69D*^, accumulate in the ER (Chow *et al.* 2016). Expression of *rpr*, *grim*, and *hid* leads to inhibition of Diap1, releasing the inhibition on initiator caspases and allows for the activation of effector caspases and downstream apoptosis. Models used in this or previous studies of retinal degeneration in the DGRP are indicated in white.

We observed substantial phenotypic variation across the DGRP for both *rpr*- and *p53*-induced apoptosis. Using genome-wide association analysis, we identified a number of modifying pathways and genes, several of which have known roles in cell death pathways, neuronal development, neuromuscular diseases, and cancer. Using systems biology approaches, we also identified Wnt signaling, mitochondrial redox homeostasis, and protein ubiquitination/degradation as possible modifiers of apoptosis. Finally, we confirmed that loss of many of these candidate modifier genes significantly alters degeneration. Our findings highlight several exciting new areas of study for apoptotic modifiers, as well as a role for stress-induced cell death in the regulation of degenerative disorders.

## METHODS

### Fly stocks and maintenance

Flies were raised at room temperature on a diet based on the Bloomington Stock Center standard medium with malt. The strains containing *GMR-GAL4* and *UAS-p53* or *GMR-rpr* on the second chromosome (*GMR>p53* and *GMR-rpr*) have been previously described (Hay *et al.* 1995; Jin *et al.* 2000). These are referred to as the apoptotic models throughout the manuscript. 204 strains from the DGRP were used for the *GMR>p53* study (Table S1) and 202 were used for the *GMR-rpr* study (Table S2). In both cases virgin females carrying one of the apoptosis models were crossed to males of the DGRP strains. F1 progeny carrying *GMR>p53* or *GMR-rpr* were collected and scored for eye size. The following RNAi and control strains are from the Bloomington Stock Center: *swim* RNAi (55961), *CG3032* RNAi (57560), *LysRS* RNAi (32967), α*Man1a* RNAi (64944), *LIMK1* RNAi (62153), *hay* RNAi (53345), *CG1907* RNAi (38998), *Sema1a* RNAi (34320), *MED16* RNAi (34012), *bru1* RNAi (44483), *CycE* RNAi (33645), *shab* RNAi (55682), *CG31559* RNAi (64671), *Cyt-c-P* RNAi (64898), *Ir40A* (57566), *sif* RNAi (61934), control *attP40* (36304), and control *attP2* (36303).

### Eye size imaging

For eye images, adult females were collected under CO_2_ anesthesia and aged to 2-7 days, then flash frozen on dry ice. Left eyes were imaged for all measurements. 10-15 eyes per strain were imaged at 3X magnification using a Leica EC3 camera. Eye area was measured in ImageJ as previously described (Chow *et al.* 2016).

### Phenotypic analysis and genome-wide association

For each DGRP line, eyes from 10-15 individual females were imaged and measured. The P-values for association of genetic background and eye size for each model were calculated using one-way ANOVA on R software. Mean eye area was used for the genome-wide association (GWA). GWA was performed as previously described (Chow *et al.* 2016). DGRP genotypes were downloaded from the website, http://dgrp.gnets.ncsu.edu/. Variants were filtered for minor allele frequency (≥ 0.05), and non-biallelic sites were removed. A total of 1,967,719 variants for *p53* and 1,962,205 variants for *rpr* were included in the analysis. Mean eye size for 2953 F1 DGRP/*GMR>p53* or 2987 DGRP/*GMR-rpr* F1 progeny were regressed on each SNP. To account for cryptic relatedness (He *et al.* 2014; Huang *et al.* 2014), GEMMA (v. 0.94) (Zhou and Stephens 2012) was used to both estimate a centered genetic relatedness matrix and perform association tests using the following linear mixed model (LMM):

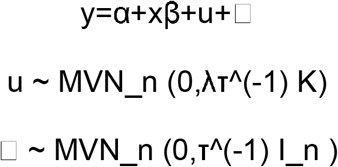

where, as described and adapted from Zhou and Stephens 2012, y is the n-vector of mean eye sizes for the n lines, α is the intercept, x is the n-vector of marker genotypes, β is the effect size of the marker. u is a n x n matrix of random effects with a multivariate normal distribution (MVN_n) that depends on λ, the ratio between the two variance components, T^(−1), the variance of residuals errors, and where the covariance matrix is informed by K, the calculated n x n marker-based relatedness matrix. K accounts for all pairwise non-random sharing of genetic material among lines. □, is a n-vector of residual errors, with a multivariate normal distribution that depends on T^(−1) and I_n, the identity matrix. Genes were identified from SNP coordinates using the BDGP R54/dm3 genome build. A SNP was assigned to a gene if it was +/-1 kb from a gene body.

### Correlation Analysis

Correlation analyses were performed to compare mean eye size in DGRP strains between *GMR>p53*, *GMR-rpr*, and *GMR>Rh1*^*G69D*^ (Chow *et al.* 2016). Statistics were calculated using a Pearson Correlation Test using R software.

### RNAi Validation

Virgin females from the apoptotic models were crossed to males carrying RNAi constructs targeting candidate modifiers of those models, and the eye size of F1 progeny expressing both the apoptotic model and the RNAi construct was measured as described above. The eyes of 10-15 females were imaged and measured. Eye size from RNAi-carrying strains were compared directly to genetically matched *attP40* or *attP2* controls using a Dunnett’s multiple comparisons test.

### Bioinformatics Analysis

Genetic polymorphisms were associated with candidate genes within 1 kb of the polymorphism. Information about candidate genes and their human orthologues was gathered from a number of databases including Flymine, Flybase, OMIM, and NCBI. Genetic interaction maps were generated using the GeneMANIA plugin on Cytoscape (version 3.6.1) (Shannon *et al.* 2003; Montojo *et al.* 2010). GSEA was run to generate a rank-list of genes based on their enrichment for significantly associated polymorphisms. For GSEA analysis, polymorphisms within 1kb of more than 1 gene were assigned to one gene based on a priority list of exon, UTR, intron, and upstream or downstream. Genes were assigned to GO categories, and calculation of enrichment score was performed as described (Subramanian *et al.* 2005). Categories with ES scores > 0 (enriched for associated genes with low p-values), gene number > 3, and p-values <0.05 were included in the final output.

## RESULTS AND DISCUSSION

### *rpr*- and *p53*-induced apoptosis is dependent on genetic background

We used the *Drosophila* eye to model apoptosis. Expression of either *p53* or *rpr* in the ommatidial array of the developing eye imaginal disc results in massive cell death and smaller, rough adult eyes (Hay *et al.* 1995; Jin *et al.* 2000). The *rpr* model is induced by direct drive of the *GMR* promoter (*GMR-rpr*) on a second chromosome balancer. The *p53* model is induced using the *GAL4/UAS* system, where *GMR-GAL4* drives expression of *UAS-p53* (*GMR>p53*). Importantly, in both of these models, adult eye size is an easily scorable, quantitative proxy for levels of apoptosis. The lines described serve as the donor strains (*GMR>p53/CyO* or *GMR-rpr,CyO/sna^sco^*) that we crossed to each DGRP strain. Females from the donor strains were crossed with males of each of 204 or 202 DGRP strains to generate F1 progeny that overexpressed *p53* or *rpr*, respectively, in the eye disc. The progeny received 50% of their genome from the maternal donor strain and 50% from the paternal DGRP strain. Therefore, we are measuring the dominant effect of the DGRP background on the *p53* or *rpr* retinal phenotype. This cross design is similar to a study of ER stress-induced degeneration (Chow *et al.* 2016) and a model of Parkinson’s Disease (Lavoy *et al.* 2018) we previously reported. We examined eye size in the F1 progeny to determine the average eye size in individual genetic backgrounds (Figure 2A-D).

**Figure 2.**
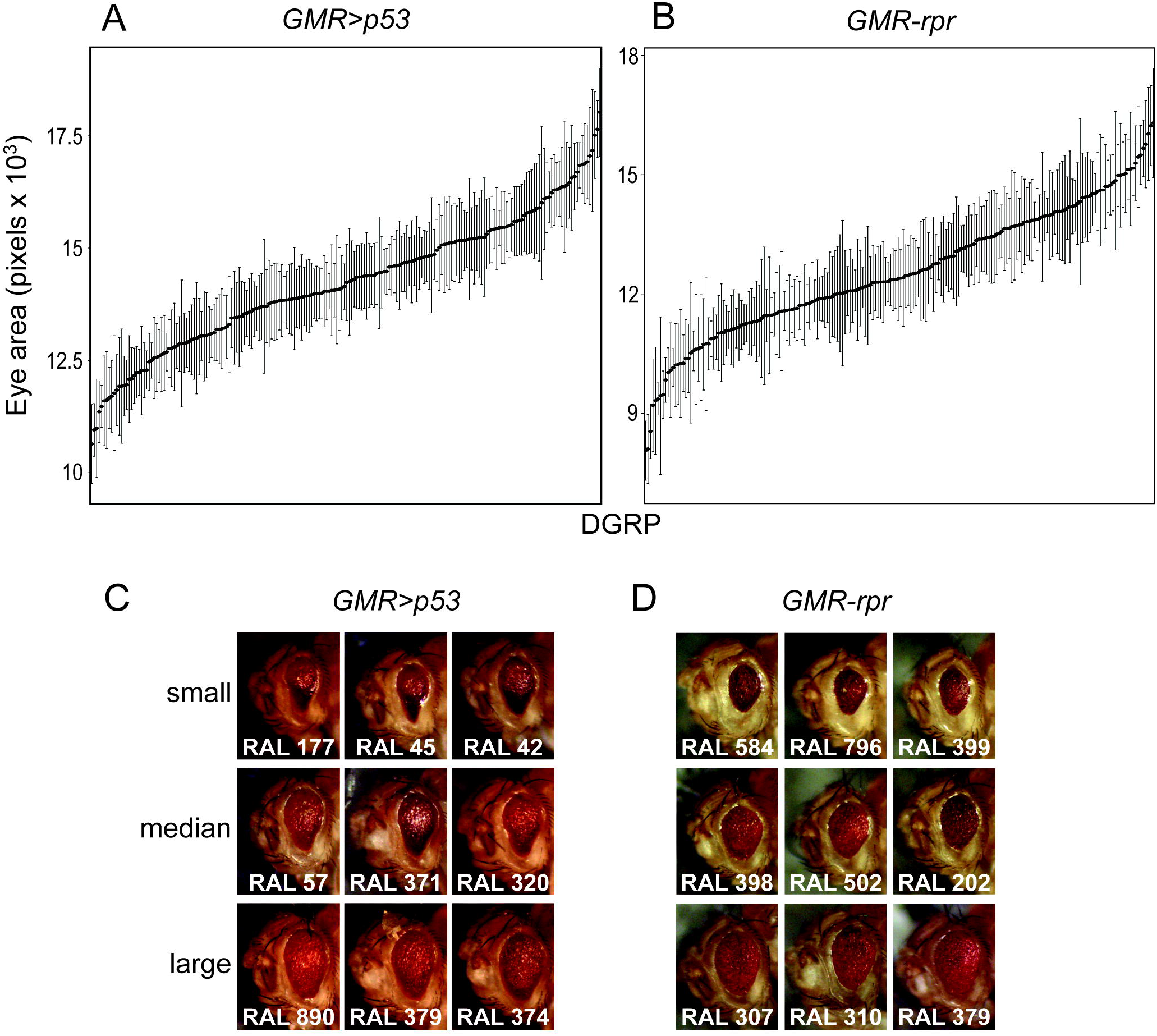
Apoptosis levels vary across genetic background in *p53* and *rpr* models of apoptosis-induced degeneration. Apoptosis induced by overexpression of *p53* (**A**) or *rpr* (**B**), as measured by adult eye size, varies across different genetic backgrounds. DGRP strains are arranged along the X-axis from smallest to largest (median eye size ± standard deviation). Representative images of *GMR>p53* eyes (**C**) or *GMR-rpr* eyes (**D**) in different DGRP backgrounds demonstrate the phenotypic variation quantified in panels **A** and **B**.

We first tested the effect of sex on apoptosis in a pilot study. We measured eye area in at least ten females and ten males from eight different DGRP strains crossed to either the *p53* or *rpr* model. Eye size is positively correlated between males and females (Figure S1A,B). Because variation is greater in females (Figure S1A,B), we elected to focus on female eye size for the remainder of our analysis.

We found a significant effect of genetic background on eye size in the *GMR>p53* model (P < 2.2 × 10^−16^) (Figure 2A,C, Table S1). Average eye size measured in pixels on ImageJ ranged from 10542 pixels (RAL812) to 17835 pixels (RAL374) (Table S1). Similarly, we found a significant effect of genetic background on eye size in the *GMR-rpr* model (P < 2.2 × 10^−16^), with median eye size ranging from 7957 pixels (RAL83) to 16884 pixels (RAL304) (Figure 2B,D, Table S2). For both the *GMR>p53* and the *GMR-rpr* models, the variation in eye size within individual DGRP strains is substantially smaller than the variation observed between DGRP strains expressing the *GMR-rpr* model (Figure 2A-B, Table S1-S2).

We noted that the range in average eye size for the *GMR-rpr* model (8927 pixels) is greater than that seen in the *GMR>p53* model (7293 pixels). This could be due to the greater involvement of *rpr* in a variety of stress-induced, *p53*-independent apoptotic pathways (Shlevkov and Morata 2012). Alternatively, it is possible that variation in *p53*-associated pathways is simply less well-tolerated than in *rpr*-associated pathways. It is also possible that the DGRP simply carries more variation affecting the *GMR-rpr* model than *GMR>p53* model.

We observed qualitative differences between the apoptotic models, with flies expressing the *GMR>p53* model displaying a teardrop-shaped eye (Figure 2C) and flies expressing the *GMR-rpr* model displaying a rounder eye (Figure 2D). These qualitative shapes were not subject to effects of genetic variation. The differences in eye shape noted between *GMR>p53* and *GMR-rpr*, however, could be indicative of differences in the mechanisms by which apoptosis and degeneration progress in these two models. Alternatively, this could be evidence of the technical differences in the two models, since *p53* is driven by the *GAL4/UAS* system and *rpr* is driven directly by the *GMR* promotor. We saw no accumulation of necrotic tissue in strains experiencing severe degeneration, nor did we note obvious differences in pigmentation (Figure 2C,D). Eyes from all strains maintained the rough-eye phenotype that is characteristic of *p53* or *rpr*-induced degeneration, indicating that while modifying variation may reduce the amount of cell death in the eye imaginal disc, it cannot fully rescue the degenerative phenotype.

### Apoptosis models correlate depending upon the pathway they activate

Because canonical p53 signaling activates the expression of *rpr*, we expected high correlation in apoptosis levels and eye size between these models (Shlevkov and Morata 2012; Mollereau and Ma 2014). Indeed, there is a significant positive correlation in eye size between DGRP strains expressing *GMR>p53* and *GMR-rpr* (r = 0.19, p = 0.0071) (Figure 3A). In a previous study, we examined the impact of genetic variation on a model of retinitis pigmentosa (RP) and ER stress-induced apoptosis (Chow *et al.* 2016). In this study, we found that the degeneration induced by overexpression of a misfolded protein (*Rh1*^*G69D*^) in the developing eye imaginal disc is modified by a number of genes involved in apoptosis (Chow *et al.* 2016). This is to be expected, as the primary cause of degeneration in this model is JNK-*hid/grim/rpr*-mediated cell death (Figure 1) (Kang *et al.* 2012). Consistent with this mechanism of *Rh1*^*G69D*^-induced degeneration, we found a significant correlation in eye size between the *Rh*1^*G69D*^ and *rpr* models (r = 0.25, p = 0.001, Figure 3B). In contrast, we see no correlation between the *Rh1*^*G69D*^ and *p53* models of apoptosis (r = 0.12, p = 0.13) (Figure 3C). These results suggest that there is shared genetic architecture between *Rh1*^*G69D*^ and *rpr*-mediated apoptosis and degeneration that is independent from that shared between *p53* and *rpr*.

**Figure 3.**
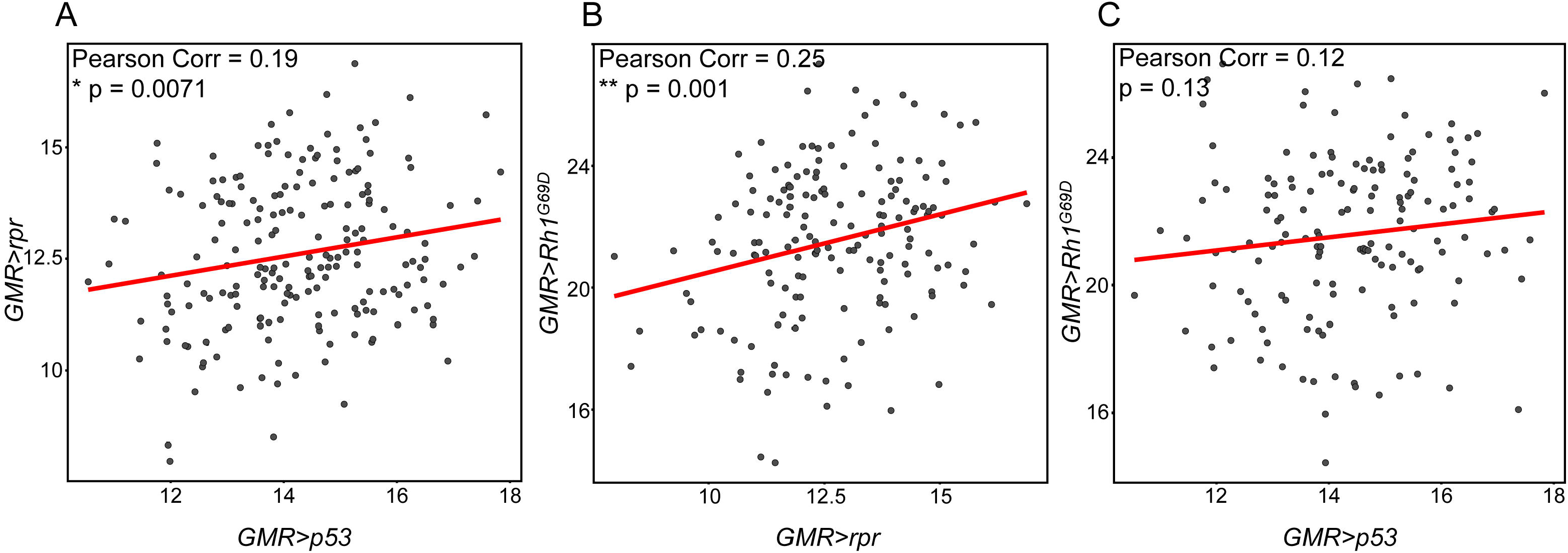
Eye size is correlated between *GMR-rpr* and both *GMR>p53* and *GMR>Rh1*^*G69D*^ models of degeneration. Correlation in mean eye size between the *GMR-rpr*, *GMR>p53*, and *GMR>Rh1*^*G69D*^ models across the DGRP. **A.** Eye size is significantly correlated in the same DGRP strains expressing *GMR-rpr* and *GMR>p53* (r = 0.19, P = 0.0071). **B.** Eye size is significantly correlated in the same DGRP strains expressing *GMR-rpr* and *GMR>Rh1*^*G69D*^ (r = 0.25, P = 0.001). **C.** Eye size is not correlated in same DGRP strains expressing *GMR>p53* and *GMR>Rh1*^*G69D*^ (r = 0.12, P = 0.13). * P < 0.05, ** P < 0.005.

### *rpr*-induced degeneration is modified by apoptosis, Wnt signaling, and mitochondrial metabolism

#### Genome-wide association analysis

To identify the genes driving this phenotypic variability, we performed a genome-wide association analysis to identify genetic polymorphisms that impact the severity of degeneration in the *GMR>p53* and *GMR-rpr* models of apoptosis. We used mean eye size as a quantitative phenotype to test for association with polymorphisms in the DGRP. Using a p-value cutoff of <1×10^−04^, we identified 128 significantly associated polymorphisms for the *GMR-rpr* model (Table S3). We only considered polymorphisms that fall within +/-1 kb of a gene. Sixteen polymorphisms lie outside of these parameters and were not considered further. Of the remaining 112 polymorphisms, ten are located in an intergenic region (+/-1kb), 14 are located in UTRs, 69 are located in introns, and 19 are located in protein-coding sequences. All 19 polymorphisms in coding regions are synonymous variants. These 112 gene-associated polymorphisms lie in 82 candidate genes (Table S3, S4). Sixty-six of the candidate genes have direct human orthologues (Table S4). A more stringent p-value cutoff (<1×10^−5^) yields only 20 polymorphisms, 16 of which lie in 14 candidate genes (12 with human orthologs) (Table S3, S4). Because the more stringent cutoff yielded few candidates, we focused the majority of our analysis on the 82 candidate genes identified at p < 1×10^−04^.

For the *GMR>p53* model, we identified 24 polymorphisms at a p-value cutoff of <1×10^−04^ (Table S5). Eight of these polymorphisms lie outside of genes and were not considered further. Of the remaining 16 polymorphisms, one is located in a UTR, 15 are located in introns, and eight are intergenic. The 16 gene-associated polymorphisms lie in 13 candidate genes (Table S5, S6). Thirteen of the associated polymorphisms have a p-value of <1×10^−05^. Five of these are intergenic, while the remaining six are in six candidate genes. Interestingly, there is no overlap between the *GMR>p53* candidate polymorphisms or genes and those identified using the *GMR-rpr* model of apoptosis (Table S3-S6). The only overlap in modifier genes is between *GMR>Rh1^G69D^* and *GMR>p53* (Table S6) (Chow *et al.* 2016). They share candidate modifier genes *CG31559*, a disulfide oxidoreductase (FlyBase Curators *et al.* 2004), and *dpr6*, a cell surface immunoglobulin involved in synapse organization (Gaudet *et al.* 2011). It is unclear what the significance of this overlap might be.

We conclude from our initial analysis that the top candidates for our models of degeneration are highly specific to the method by which we induce that degeneration. Because there are so few significant associations for the *GMR>p53* model of apoptosis, and even fewer that are in close proximity to a candidate gene, we elected to focus the remaining analysis on the *GMR-rpr* model.

#### Modifier genes

Because the *rpr* model directly induces apoptosis, we expected to see apoptotic functions for many of the candidate genes identified in our GWAS. The top hit was the gene *echinus* (*ec*), a ubiquitin specific protease (USP) orthologous to human *USP53* and *USP54* (Table S4). We identified nine intronic SNPs in *ec* through our association analysis. Previous studies show that loss of *ec* in the developing eye results in a mild rough eye phenotype, albeit a much less dramatic one than that seen upon overexpression of *rpr* (Wolff and Ready 1991; Copeland *et al.* 2007). While this previous study reported no genetic interaction between *ec* and *rpr*, this was assessed based on qualitative changes as opposed to quantitative differences in eye size (Copeland *et al.* 2007). Our GWAS data suggests that such a genetic interaction may play an important role in *rpr*-induced degeneration.

*Ec* is one of several apoptotic genes identified in this analysis. In fact, 16/82 (~20%) of the candidate genes have known functions in apoptosis-related pathways, all of which have conserved human orthologues (Table S4). One of these is *Diap2*, a *Drosophila* paralog of *Diap1* (human orthologs: *BIRC2* and *BIRC3*) (Hay *et al.* 1995). The Diap proteins normally inhibit caspase activation and prevent apoptosis. Expression of the rpr/grim/hid proteins inhibits Diap1 and Diap2, allowing apoptosis to proceed. Increased expression or activity of Diap2 reduces the impact of *rpr* overexpression, thereby reducing apoptosis (Hay *et al.* 1995). Conversely, reduced expression of *Diap2* may not have a strong impact on *rpr*-associated degeneration, as *Diap1* is the major functional paralog in this pathway. The identification of a gene directly involved in the *rpr* pathway demonstrates the efficacy of our GWAS.

Two candidates, *hay* and *Xpd* (*ERCC3* and *ERCC2*) (Table S4), have human orthologs mutated in Xeroderma pigmentosum, an inherited genetic condition where defects in DNA excision repair result in melanomas and eventually death (Kraemer and DiGiovanna 2016). These are subunits of the TFIIH helicase complex that are involved in excision repair after UV damage (Koken *et al.* 1992; Mounkes *et al.* 1992; Reynaud *et al.* 1999). Besides *hay* and *Xpd*, we identified 4 additional genes whose human orthologs are directly involved in cancer: *DIP-iota* (*OPCML*), *Fum4* (*FH*), *CG8405* (*TMEM259*), and *CG15529* (*BLNK*). Mutations in these genes have been associated with ovarian cancer (*OPCML*) (Sellar *et al.* 2003), renal cancer (*FH*) (The Multiple Leiomyoma Consortium 2002; Pollard *et al.* 2005), and various carcinomas (*TMEM259*) (Chen *et al.* 2005). The roles of these genes in cancer are likely due to functions in apoptotic initiation or cell cycle regulation. Other candidates are activated downstream of p53, such as *CG44153* (*ADGRB3*) and *stac* (*BAIAP3*) (Shiratsuchi *et al.* 1997, 1998). This suggests that feedback signaling through p53 can increase *rpr*-induced apoptosis and degeneration.

24/82 candidate genes (~30%) are involved in neuronal function or implicated in neurological disease. Twenty-three have conserved human orthologues (Table S4). Human orthologs of *Form3* (*INF2*) and *LysRS* (*KARS*) can both be mutated in different forms of the degenerative peripheral neuropathy Charcot-Marie-Tooth disease (McLaughlin *et al.* 2010; Boyer *et al.* 2011), while *Shawl* (*KCNC3*) and *CG7741* (*CWF19L1*) are associated with spinocerebellar ataxia (Waters *et al.* 2006; Burns *et al.* 2014). Mutation in the Rab3-interacting scaffold protein encoded by *Rim* (*RIMS1*) can cause a retinal degenerative disease that is similar to retinitis pigmentosa (Johnson *et al.* 2003), which was the focus of the *Rh1*^*G69D*^ study (Chow *et al.* 2016). Identification of genes with roles in different neuronal and muscular degenerative diseases suggests that these modifiers could be important in a variety of apoptosis-associated diseases.

#### Network analysis

To understand if there are functional relationships between *GMR-rpr* modifiers, we examined interactions among the 82 candidate genes. Genetic, physical, and predicted interactions were compiled and visualized using Cytoscape software (Shannon *et al.* 2003; Montojo *et al.* 2010). Fourteen of the 82 candidate genes were found as nodes in these interaction networks, as was *rpr* itself (Figure 4A). We identified several interesting clusters of candidate genes, including those with functions in apoptosis, development, and protein ubiquitination.

**Figure 4.**
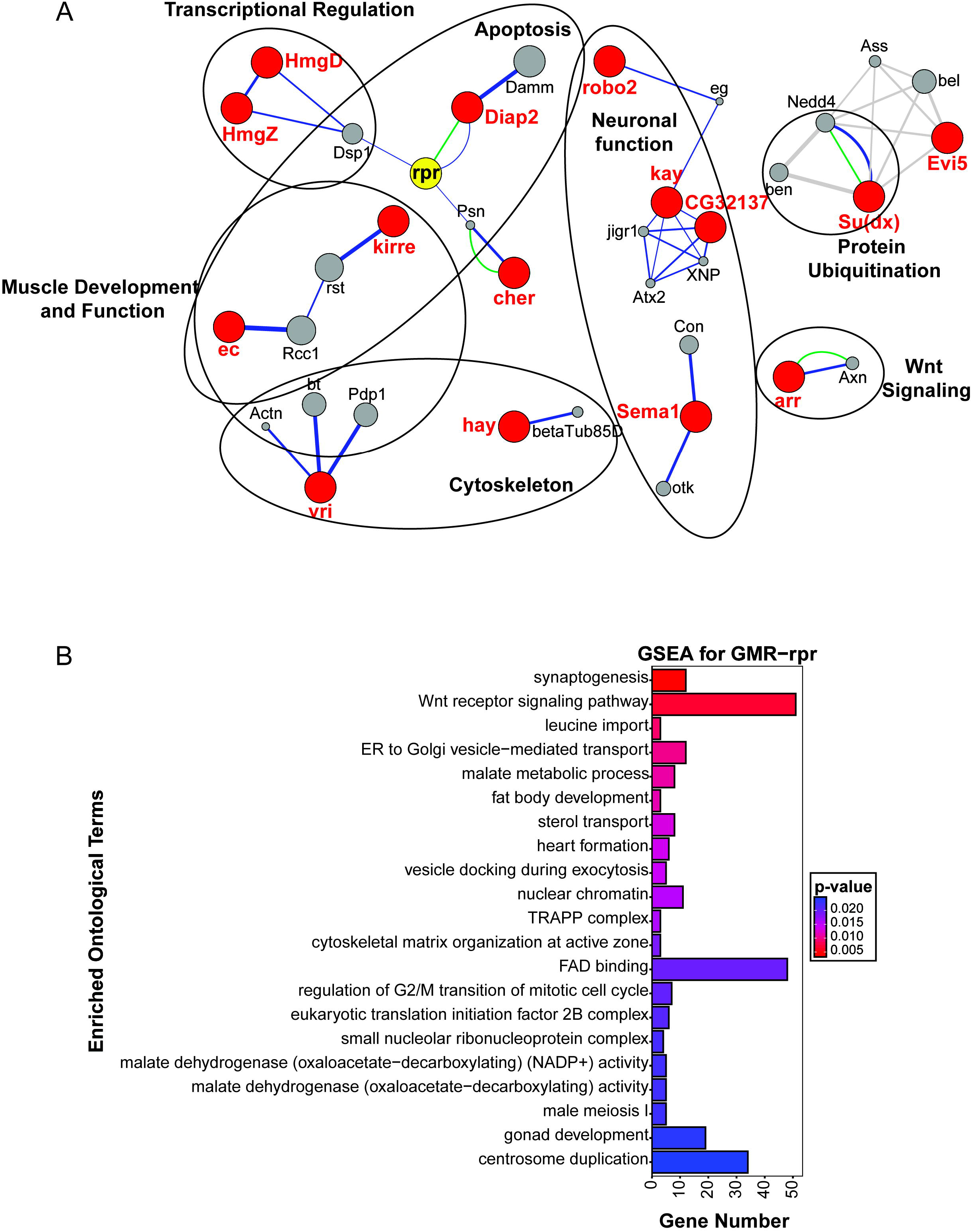
*rpr* modifiers are enriched for neuronal function, Wnt signaling, and metabolic pathways. **A**. *rpr* modifier network, as plotted by the GeneMANIA plugin in Cytoscape (Shannon *et al.* 2003; Montojo *et al.* 2010). Significant candidate modifiers are indicated in red, with physical interactions shown in green, genetic interactions shown in blue, and predicted interactions shown in gray. Circles represent groups with a functional identifier. **B.** Top 20 significant ontological categories as identified by GSEA. Categories are arranged from most significant on top to least significant along the y-axis. P-values are indicated by red-to-blue gradient, with red the lowest p-values and blue the highest P-values. Gene number identified in each category is indicated along the y-axis.

As expected, given the large number of candidates with apoptotic roles, we found an apoptosis cluster of interactions between modifiers with functions associated with cell cycle regulation and cell death (Figure 4A). A number of these genes, including *Diap2* and *cher* (*FLNA*), have either direct or indirect interactions with *rpr* itself. As noted above, *Diap2* interacts both physically and genetically with *rpr* (Figure 1) (Hay *et al.* 1995). *cher* shows indirect genetic interactions with *rpr* through its physical association with the presenillin (*psn*) protein (Guo *et al.* 2000) (Figure 4A). This interaction is conserved in humans and is specifically associated with Alzheimer’s Disease (Lu *et al.* 2010).

We also observed a cluster composed of regulators of developmental apoptosis in our network, including the *ec* protease (Copeland *et al.* 2007) and the neuronal cell adhesion protein encoded by *kirre* (*KIRREL3*) (Bao *et al.* 2010) (Figure 4A). Indirect genetic interactions were identified between these genes, which are commonly involved in development of the *Drosophila* eye imaginal disc and accompanying regulated apoptosis. The chromatin-binding HmgD/Z (*HMGB2*) proteins are expressed at high levels in the larval CNS, suggesting that they are important for the developmental regulation of neuronal gene expression (Churchill *et al.* 1995; Gaudet *et al.* 2011; Brown *et al.* 2014). They indirectly interact with *rpr* through the closely related *dsp1*, which encodes another paralog of human *HMGB2* (Figure 4A). Dsp1 recruits members of the repressive polycomb complex to chromatin. It is possible that these genetic interactions indicate a role for the HMGB2 proteins in regulating *rpr* expression and, as a result, developmental regulation of cell death and tissue turnover. Our apoptotic model is expressed in a developmental tissue, suggesting that some of the variation in eye size observed across the DGRP could be due to changes in the response of developmental processes to the abnormal activation of apoptosis. Such regulators of developmental apoptosis could be excellent candidates for therapeutic targeting in degenerative diseases.

We also identified a number of predicted interactions in a cluster of modifier genes involved in protein ubiquitination (Figure 4A). Among the top candidate genes are *ec*, *Diap2*, *Su(dx)* (*ITCH*), and *Roc2* (*RNF7*), all of which have important roles in protein degradation through ubiquitination and the proteasome degradation pathway. *Su(dx)*, like *Diap2*, encodes a ubiquitin ligase (Gaudet *et al.* 2011). Our network analysis highlights a predicted interaction between *Su(dx)* and the Rab GTPase-interacting protein *Evi5*, another candidate gene (Laflamme *et al.* 2012) (Figure 4A). This regulator of vesicular fusion is predicted to interact with a number of additional ubiquitin ligases as well (Figure 4A). Degradation of proteins through the proteasome is an important mechanism for maintaining cellular homeostasis under a variety of cellular stresses (Sano and Reed 2013). Altered regulation of E3 ligases, which determine the identity and specificity of proteins for degradation (Ester Morreale and Walden 2016), could tip the balance of cells experiencing apoptotic stress toward or away from cell death.

#### Gene set enrichment analysis

Thus far, we have focused our analysis on rank-order candidate modifiers identified in our GWAS. While this provides many new avenues for future analysis, it ignores the majority of the association data. We therefore performed gene set enrichment analysis (GSEA), using all GWAS variant data and their associated P-values. The gene nearest to each variant was assigned the variant’s P-value and used as GSEA input, using the method described (Subramanian *et al.* 2005). Given a defined set of genes annotated with a certain GO function, GSEA determines whether the members of that set are randomly distributed throughout the ranked list or if they are found primarily at the top or bottom of that list. GO categories enriched at the top of the list functionally describe the phenotype of the gene set. While traditional GO analysis uses a set of genes based on a P-value cutoff, GSEA examines the entire gene set (Dyer *et al.* 2008). GSEA identified 62 significantly associated gene sets (≥ 3 genes) at a p-value of <0.05 (Table S7). The top gene set was synaptogenesis (GO:0007416, P = 3.7 × 10^−3^) and includes *Sema1a* (*SEMA6A*), a conserved semaphorin-binding protein involved in axon guidance (Ayoob *et al.* 2006; Gaudet *et al.* 2011) and one of the top modifier candidates based on individual polymorphism analysis (Figure 4B, Table S7). Other genes in this category include those involved in synapse formation and organization, suggesting that regulating neuronal connectivity and synapse choice could play a role in the decision to apoptose or to survive.

The second most significantly enriched category was Wnt signaling (GO:0016055, P = 6.7 × 10^−3^), consisting of 51 enriched genes from our *GMR-rpr* analysis (Figure 4B, Table S7). One of these, *arr*, is also a candidate modifier gene (Table S4,S7). *arr* is a *Drosophila* orthologue of the genes encoding the co-receptors *LRP5/6* in canonical Wnt signaling (Rives *et al.* 2006). The second most significant single candidate gene in the GWA is *swim* (*TINAGL1*/*TINAG*), a secreted cysteine protease capable of binding the wingless (wg) ligand and enhancing its spread and signaling capabilities (Mulligan *et al.* 2012). Also enriched for significant polymorphisms are four frizzled paralogs (Wnt receptors) and six paralogs of the Wnt ligand (Table S7). Other integral components of the canonical Wnt pathway, such as *disheveled*, *axin*, and *CKI*α, are enriched for associated polymorphisms, as are several peripheral and non-canonical regulators of Wnt signaling (Table S7). This striking association is reinforced by previous studies that have linked Wnt signaling with either the promotion or restraint of cell death (Pećina-Slaus 2010). Non-canonical Wnt signaling can activate JNK or calcium release from the ER, both of which can alter the decision to initiate apoptosis (Rasmussen *et al.* 2018). It will be interesting to investigate Wnt signaling collectively as well as with individual candidates to determine how different branches of the pathway impact degenerative diseases.

GSEA also identified a number of genes and pathways involved in mitochondrial homeostasis and metabolism (Figure 4B), including malate metabolic processes (seven genes, GO:0006108, P = 0.011). These genes encode for malate dehydrogenase enzymes, six of which are localized to the mitochondrion (Figure 4B, Table S7). Malate dehydrogenase catalyzes the oxidation of malate to oxaloacetate in the last step of the TCA cycle prior to the entrance of acetyl-CoA (Minárik *et al.* 2002). The presence of so many paralogs of this enzyme suggests that mitochondrial metabolism, and in particular the mitochondrial redox state, is a major regulator of apoptosis. Supporting this, one of the top candidates, Fum4 (*FH*), is also an essential enzyme in the TCA cycle (Table S4). The GSEA further supports this finding, as FAD binding is also enriched (48 genes, GO:0050660, P = 0.020) (Figure 4B). A primary function for these 48 enriched genes is the maintenance of redox homeostasis, 16 of which localize to the mitochondria. Another of these genes, the apoptosis-inducing factor *AIF*, is activated independently from caspases by mitochondrial stress and is released into the cytoplasm, travels to the nucleus, and initiates the chromatin condensation and DNA fragmentation that immediately precedes cell death (Elmore 2007).

More generally, redox homeostasis in other cellular compartments is also implicated by GSEA (Table S7, Figure 4B). Three paralogs of aldehyde oxidase (*Aox*) and the NAD(P)H oxidoreductase *Duox* (*DUOX1*) are enriched for associated polymorphisms; these oxidase enzymes are essential for maintaining an appropriate balance of reactive oxygen species in the cytoplasm. We identified four paralogs of acyl-coA oxidase (*Acox*), which is involved in the β-oxidation of very long chain fatty acids in the peroxisome, and an additional 4 genes involved in mitochondrial β-oxidation: *wal* (*ETFA*), *Mcad* (*ACADM*), *CG4860* (*ACADS*), and *CG7461* (*ACADVL*).

The involvement of enzymes regulating redox homeostasis, and more specifically redox homeostasis in the mitochondria, is consistent with *rpr*-induced apoptosis. Both caspase-dependent and caspase-independent apoptotic pathways can be activated downstream of mitochondrial stress (Elmore 2007; Rasmussen *et al.* 2018). Increasing the permeability of the mitochondrial membrane is sufficient to ensure activation of the apoptosome through the release of cytochrome-C (Elmore 2007). This, along with expression of the mitochondria-associated IAP inhibitors rpr/grim/hid, activates the caspase cascade (Sandu *et al.* 2010). Damage to the mitochondria that increases permeability, such as through redox stress, is itself sufficient to activate apoptosis in a caspase-independent manner through the release of AIF (Elmore 2007).

Other metabolic processes such as sterol transport (GO:0015918, P = 0.013), leucine import (GO:0060356, P = 8.9 × 10^−3^), and fat body development (GO:0007503, P = 0.011) are enriched in the GSEA (Table S7, Figure 4B). Disruption of metabolic processes has long been known to induce oxidative and ER stress, both of which are capable of activating apoptosis through JNK/*grm-rpr-hid* signaling cascades or directly through mitochondrial stress (Kanda and Miura 2004). It will be interesting to explore how these metabolic processes alter apoptosis not only in this model of retinal degeneration, but in physiologically relevant cell types and tissues, such as the midgut, fat body, and insulin-producing cells.

The enrichment of multiple metabolic categories suggests that the impact of cellular and mitochondrial metabolism on redox homeostasis could play a major role in *rpr*-induced degeneration. We hypothesize that these regulators of mitochondrial redox state and metabolism are directly and indirectly influencing the activation of mitochondrial proteins involved in the final decision to undergo apoptosis. Our GSEA emphasizes the importance of exploring not just individually associated genes but also their functional pathways and partners when identifying genetic modifiers of disease.

### Functional analysis of candidate modifiers of apoptosis

To confirm the roles of our candidate genes in regulating apoptosis, we elected to test the impact of loss of modifier expression for nine of the most significant rank-ordered candidate genes. We crossed RNAi targeting each of these modifiers into the *GMR-rpr* or *GMR>p53* line, and then measured the eye area in offspring carrying both the RNAi construct and the apoptosis model (Figure 5, Figure S2). Eye area was quantified and compared to a genetically matched control expressing only the apoptosis model (Figure S3A). Due to a lack of highly significant candidate modifiers of *p53*-induced apoptosis, we focus our analysis here on the *rpr* modifiers. Knockdown of either *LIMK1* (*LIMK1*) (16183 ± 875 pixels, N = 15) or *swim* expression (15518 ± 2418 pixels, N = 14) resulted in enhancement of the apoptosis phenotype, showing a significant decrease in eye size compared to controls expressing only *GMR-rpr* (17534 ± 1098 pixels, N = 11) (Figure 5). Knockdown of *sema1a* (18990 ± 746 pixels, N = 15), *MED16* (*MED16*) (20323 ± 622 pixels, N = 15), or *hay* (20240 ± 617 pixels, N = 14) resulted in a partial rescue, with a significant increase in eye size compared to controls expressing *GMR-rpr* (Figure 5). No significant change in eye size was observed upon knockdown of *CG3032* (*GZF1*) (18525 ± 449 pixels, N = 12), *LysRS* (*KARS*) (17879 ± 1834 pixels, N = 12), α*Man-1A* (*MAN1A2*) (17842 ± 763 pixels, N = 15), or *CG1907* (*SLC25A11*) (18755 ± 787 pixels, N = 13) (Figure 5). No phenotype was observed for knockdown of any of the modifiers under non-apoptosis conditions (Figure S3B,C). These results demonstrate that many of the top GWA candidate modifiers are capable of modifying the apoptotic phenotypes associated with the *GMR-rpr* model of degeneration. In the future we will also examine the impact of overexpression of candidate genes on the GMR-rpr model of apoptosis, as some candidate genes may exert a stronger influence under conditions of gain rather than loss of function.

**Figure 5.**
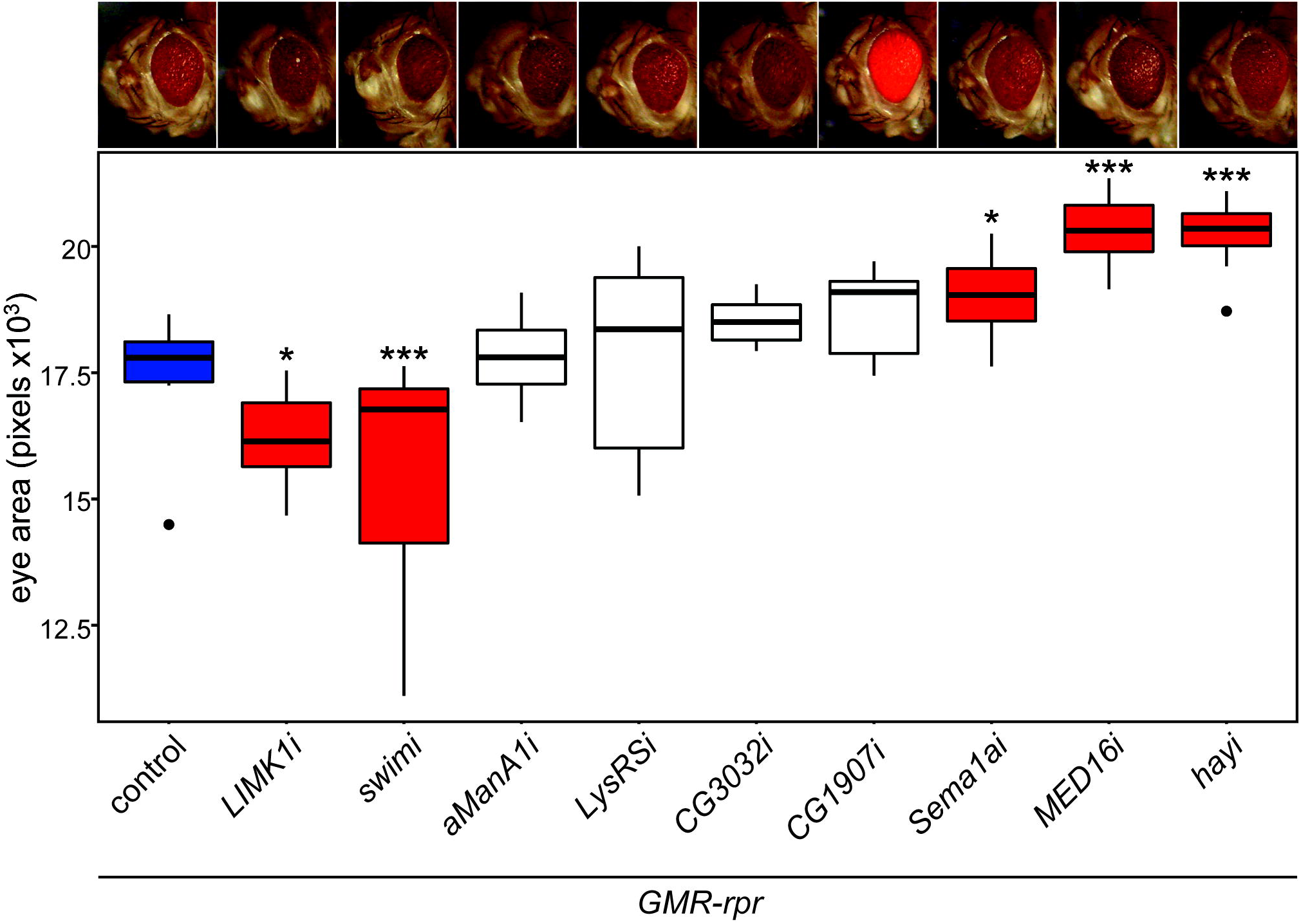
Knockdown of candidate *rpr* modifiers significantly alters apoptosis-induced degeneration. RNAi against candidate modifiers was expressed under the control of *GMR-GAL4* in the *GMR-rpr* model. The genetically matched attP2 line was crossed into the *GMR-rpr* line as a control (blue). Eye size in pixels was quantified for N = 11-15 flies per strain and plotted with the 25^th^-75^th^ percentile of measurements in the central box. Measurements lying outside of 1.5 x interquartile range are indicated as points. Representative images of each line are found above the data for that line. Knockdown of *LIMK1* or *swim* significantly reduces eye size in the *GMR-rpr* model of degeneration compared to controls. Loss of *Sema1a*, *MED16*, or *hay* results in a significant increase in eye size compared to controls. Loss of *CG1907* does not significantly alter eye size, but changes in pigmentation are similar in the presence or absence of GMR-rpr (SUPP FIG). Loss of α*ManA1*, *LysRS*, or *CG3032* do not produce a significant effect. RNAi lines with significant changes in eye size are indicated in red, while those that are not significantly changed are indicated in white. * P < 0.05, *** P < 0.0005.

## CONCLUSIONS

The primary goal of this study was to identify candidate genes and pathways that modify apoptosis and degenerative processes. Apoptosis is a primary cause of disease in a multitude of degenerative disorders (Mattson 2000). It is also a commonly targeted pathway for cancer therapies (Ouyang *et al.* 2012). These and other diseases are subject to a large degree of phenotypic heterogeneity due to inter-individual differences in genetic background among patients (Queitsch *et al.* 2012; Chow 2016). Understanding how genetic diversity in the population impacts apoptosis could therefore lead to identification of prognostic predictors in the diagnosis of disease and of new therapeutic targets. The modifiers identified here inform our understanding of cell death regulation and could serve as therapeutic targets in a variety of apoptosis-related disorders.

This study demonstrates the use of the DGRP to identify modifiers that are extremely specific to the disease model being tested. Even though the end point of degeneration is superficially the same for *rpr* and *p53* models, candidate modifier genes are unique to each model. Our methods clearly identify genetic variation associated with specific disease mechanisms, and not simply genes involved in general cell health and survival. We believe the modifying genes and pathways discovered and discussed here are excellent candidates in the treatment and understanding of apoptosis-related disorders. With further analysis, we can characterize the roles these modifiers play in degeneration and their specific functions across tissues and disease models. The genes and pathways identified study have tremendous value as possible therapeutic targets or prognostic markers of disease.

## ACKNOWLEDGEMENTS

We thank Andre Cruz and Demi Perez for their contributions to this project. This research was supported by an NIH/NIGMS R35 award (1R35GM124780) and a Glenn Award from the Glenn Foundation for Medical Research to CYC. CYC is the Mario R. Capecchi Endowed Chair in Genetics. RASP is supported by a NIDDK T32 fellowship (5T32DK110966). KGO is supported by the Genetics T32 Fellowship from the University of Utah. EO and KS were supported by the Undergraduate Research Opportunities Program (UROP) through the University of Utah Office of Undergraduate Research. AGG is supported by an NIH/NIAID R21 award (R21AI128103).

