## Supplemental Figures 1-3 for "Natural genetic variation screen in *Drosophila* identifies Wnt signaling, mitochondrial metabolism, and redox homeostasis genes as modifiers of apoptosis"

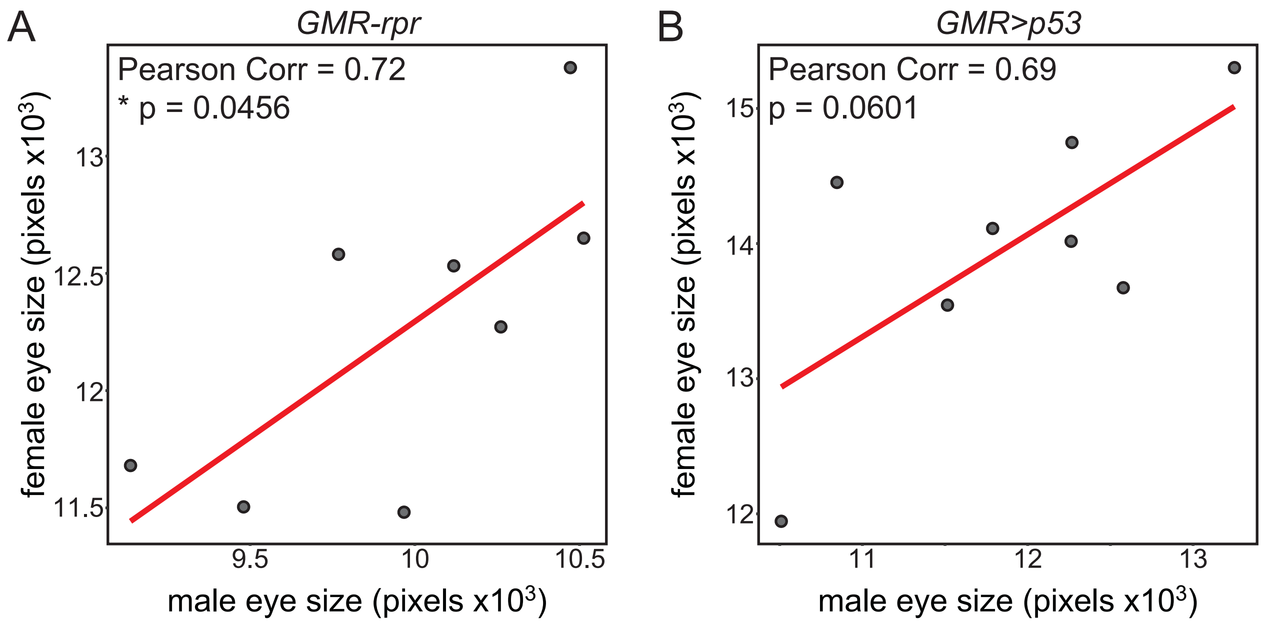


**Figure S1. Eye size varies similarly across genetic background between males and females.**

**A.** Eye size in DGRP strains expressing *GMR-rpr* show a significant positive correlation between males and females of those strains (r = 0.72, P = 0.0456). **B.** Eye size in DGRP strains expressing *GMR>p53* show a positive, but non-significant correlation between males and females (r = 0.69, P = 0.0601). Eye size in pixels x $10^3$ from select DGRP lines was compared between males (X-axis) and females (Y-axis) expressing the *GMR-rpr* or *GMR>p53* model. Correlations were calculated using a Pearson correlation test. Individual data points represent one DGRP strain. Correlation slope is indicated by the red line. * $P < 0.05$.

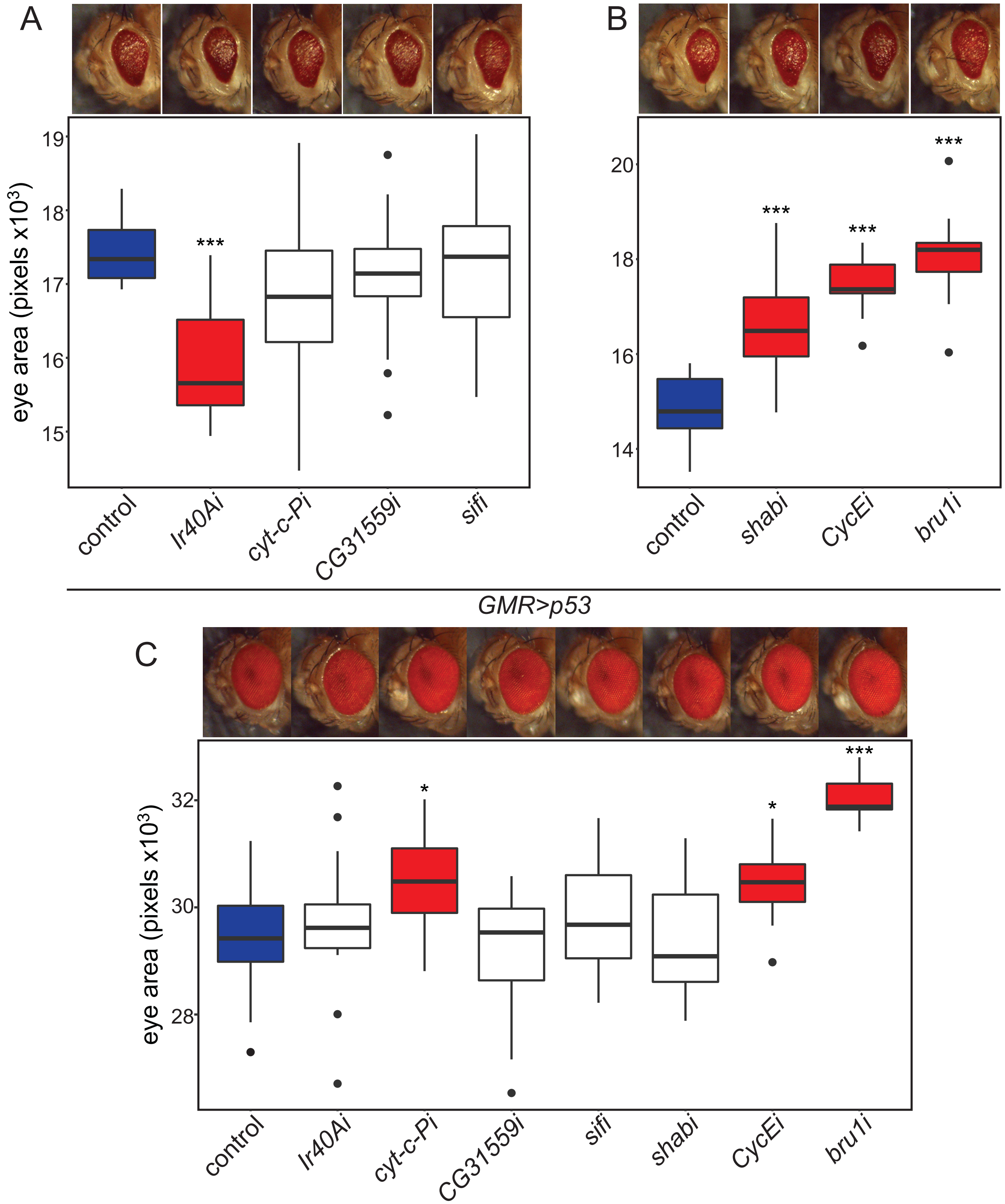


**Supplemental Figure 2. Loss of candidate *p53* modifiers significantly alters eye size in the presence and absence of p53 overexpression.**

RNAi against candidate modifiers was expressed under the control of *GMR-GAL4* in the *GMR>p53* model. The genetically matched *attP40* **(A)** or *attP2* **(B)** lines were crossed into the *GMR>p53* line controls (blue). Eye size in pixels was quantified for N = 9-15 flies per strain. Representative images of each line are found above the data for that line. **A.** Reducing *Ir40A* expression through RNAi significantly reduces eye size in the *GMR>p53* model of degeneration compared to the *attP40* control. Loss of *cyt-c-P*, *CG31559*, or *sif* does not produce a significant effect. **B.** Reducing *shab*, *CycE*, and *bru1* expression through RNAi significantly increases eye size in the *GMR>p53* model of degeneration compared to the *attP2* control. **C.** RNAi against candidate modifiers listed was expressed under the control of *GMR-GAL4* in a wild-type background. The genetically matched *attP2* line was crossed into the *GMR-GAL4* line as a control (blue). Eye size in pixels was quantified for N = 12-15 flies per strain. Representative images of each line are found above the data for that line. Reducing *cyt-c-P*, *CycE*, and *bru1* expression using the *GMR-GAL4* driver alone significantly increases eye size compared to the control strain. Knockdown of *Ir40A*, *CG31559*, *sif*, or *shabi* did not produce a significant effect. Control lines are indicated in blue. RNAi lines with significant changes in eye size are indicated in red, while those that are not significantly changed are indicated in white. * $P < 0.05$, *** $P < 0.0005$.

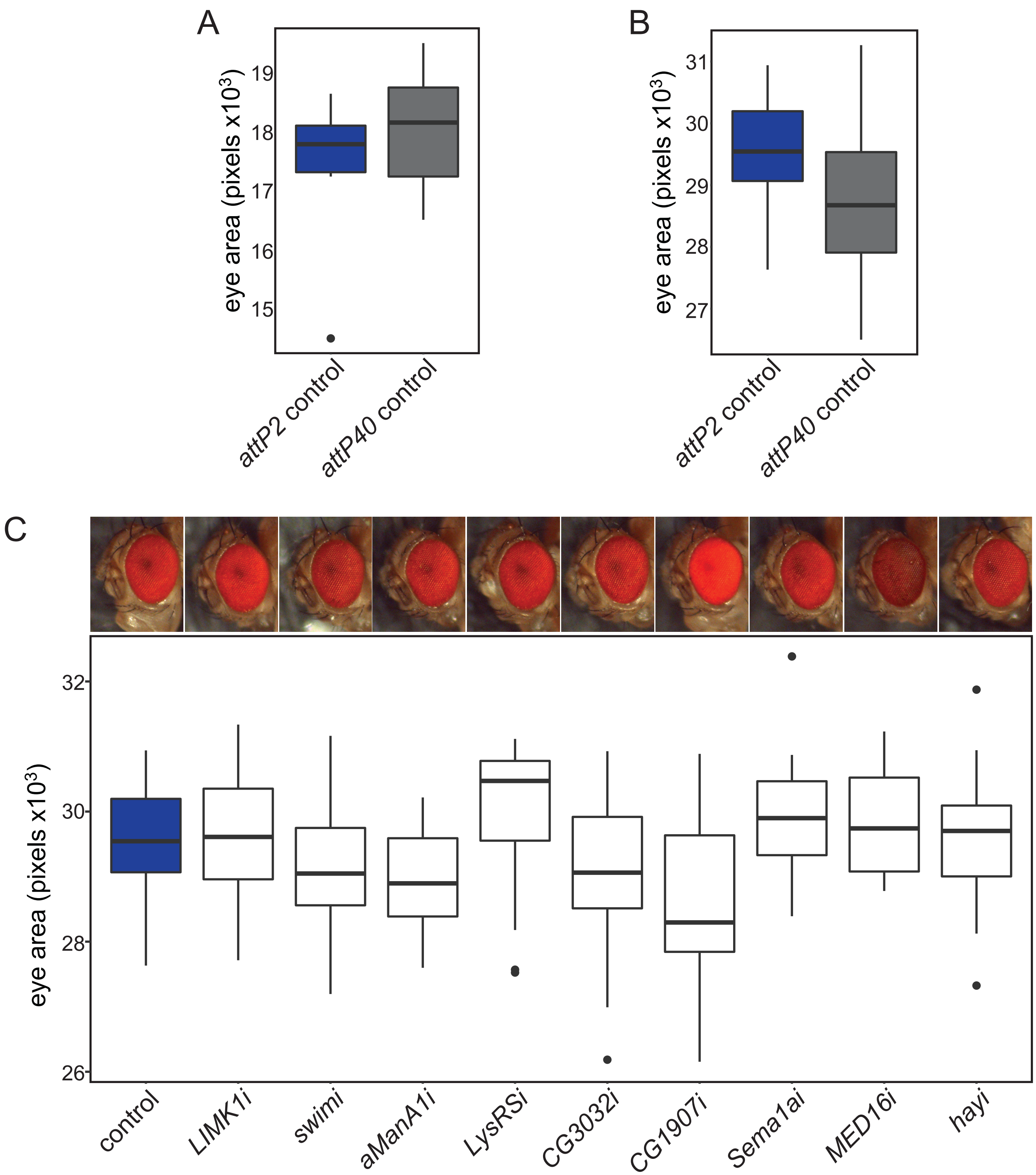


**Supplemental Figure 3. *GMR-rpr* and *GMR-GAL4* controls for candidate *rpr* modifier genes. A.** Degeneration induced by *GMR-rpr* does not significantly differ when *rpr* is expressed in the *attP2* (blue) (17534 ± 1098 pixels, N = 11) and *attP40* (gray) (18019 ± 934 pixels, N = 11) control genetic backgrounds. **B.** Eye size does not significantly differ when *GMR-GAL4* is expressed alone in the *attP2* (blue) (29472 ± 964 pixels, N = 15) and *attP40* (gray) (28800 ± 1161 pixels, N = 15) control genetic backgrounds. **C.** RNAi against candidate modifiers was expressed under the control of *GMR-GAL4* in a wild-type background. The genetically matched *attP2* line was crossed into the *GMR-GAL4* line as a control (blue). Eye size in pixels was quantified for N = 13-15 flies per strain. Representative images of each line are found above the data for that line. Knockdown of candidate gene expression using the *GMR-GAL4* driver alone does not significantly alter eye size. Qualitative changes in pigmentation were noted for *CG1907* and *MED16*. Control lines are indicated in blue, and RNAi lines that are not significantly changed are indicated in white.
